# Development of antisense tools to study *Bodo saltans* and its intracellular symbiont

**DOI:** 10.1101/2024.07.27.605423

**Authors:** Mastaneh Ahrar, Lorna Glenn, Marie Held, Andrew Jackson, Krzysztof Kus, Gregory D.D. Hurst, Ewa Chrostek

## Abstract

Obligate symbioses are common in nature and present a particular challenge for functional genetic analysis. In many cases, the host is a non-model species with poor tools for genetic manipulation and the symbiont cannot be cultured or its gene expression manipulated to investigate function. Here we investigated the potential for using antisense inhibition to analyse host and symbiont gene function within an obligate aquatic symbiosis. We focused on the kinetoplastid host *Bodo saltans* and its bacterial symbiont, C*andidatus* Bodocaedibacter vickermanii, a member of *Rickettsiales*. We conclude that antisense inhibition is not feasible in the *B. saltans* and its symbiont, as the holobiont feeds on the antisense molecules – and increases in numbers – upon treatment with the antisense construct. Although our approach has proven unsuccessful, we have developed an array of protocols which can be used to study the biology of this microeukaryote and its microbial associates.

## Introduction

Dependent symbioses – where a host requires a symbiont for function – are common in nature. They also present therapeutic opportunities in some cases, as in filarial diseases where targeting of the symbiont enables sterilization of the host and ultimately a novel treatment strategy (1, 2). More widely, many blood feeding vectors and agriculturally important phloem feeding insects rely on symbiont presence, such that understanding the basis of dependence is important in health and food security (3–9). However, the symbionts (and sometimes the host) are commonly refractory to functional analysis, inhibiting our capacity to understand the interplays resulting in dependence.

Microeukaryotic hosts present an opportunity for understanding symbiotic relationships. A broad range of symbiotic microbes inhabit single cell eukaryotes, with diverse impacts on their host. For example, in the *Paramecium – Chlorella* symbiosis the algal endosymbiont provides photosynthetic capability to the microeukaryote host (10). Additionally, the stability of this symbiosis is ensured by the “penalty system” acting on the host upon the death of the symbiont (11). Killing of the symbiont releases its mRNA with a high level of sequence identity to host transcripts. These mRNAs are then processed by the host RNAi machinery resulting in knockdown of endogenous host gene expression which is detrimental to the host. Therefore, RNA-RNA interactions can be crucial in maintaining symbiosis (11). The simplicity of microeukaryote culture, their susceptibility to RNA-level manipulation of gene expression, and possibility of delivery of small molecules to cells through the culture medium makes them a potentially useful tool in enabling functional analysis. However, development of these tools for aquatic symbioses is in its infancy.

To this end, we wished to establish tools for functional analysis in the interaction between the free-living flagellated kinetoplastid *Bodo saltans* and its intracellular bacterium, C*andidatus* Bodocaedibacter vickermanii (Cbv), a member of *Rickettsiales* (12). *Bodo saltans* is a unicellular eukaryote hosting Cbv (12). It is a heterotroph that feeds on extracellular bacteria, and is found in freshwater and marine environments (13). Cbv appears to be a permanent inhabitant of *B. saltans* cells (12). An obligate symbiosis between *B. saltans* and Cbv has been postulated, based on the observation that antibiotic clearance of the bacterium kills the eukaryote as well (12). Genomic analysis of Cbv revealed the presence of the Bacterial Polymorphic Toxin Systems (PTS) with putative role in symbiosis maintenance. It has been hypothesized that the removal of the symbiont with antibiotics would stop the production of the antitoxin and render *B. saltans* cells defenceless in the presence of a longer lived bacterial toxin.

In order to study *B. saltans* - Cbv symbiosis in more detail, we need tools for simultaneous manipulation of gene expression in host and symbiont. Antisense inhibition is a good candidate for a universal protocol working across different kingdoms. It has been achieved multiple times in trypanosomes closely related to *B. saltans*. In *Trypanosoma cruzi*, Surface Glycoprotein gp90 was blocked by the antisense approach, using a phosphorothioate oligonucleotide based on a sequence of the *gp90* coding strand (14). Similarly, expression of Calcineurin B of this parasite has also been knocked down to show that this protein is involved in cell invasion (15). Finally, *T. cruzi*, Inositol 1,4,5-Trisphosphate Receptor has also been inhibited by the antisense oligonucleotide treatment (16).

Antisense inhibition can also be applied to bacteria (17, 18) including *Rickettsia* and *Ehrlichia* (19, 20), which are closely related to Cbv. Synthetic peptide nucleic acid molecules (PNAs) targeting mRNA for rOmpB (Rickettsial outer-membrane protein B) in *R. typhi* and rickA (Arp2/3 complex activator) in *R. montanensis* have successfully induced antisense inhibition of these genes, confirming their importance for infection (19). Additionally, PNAs have been used to knock down Ehrlichial translocated factor-1 (Etf-1), which induces Rab5-regulated autophagy to provide host cytosolic nutrients required for *E. chaffeensis* proliferation. This knockdown reduced the bacteria’s ability to infect host cells (20). Importantly, intracellular localization of *Rickettsiales* often requires carriers enabling antisense molecules movement across membranes. Molecular carriers have been successfully employed to deliver specific and lasting antisense inhibition to intracellular *Chlamydia* (21) and DNA to another Cbv relative – *Anaplasma* (22).

As the development of genetic tools for the host *B. saltans* is beginning to gain traction (23, 24), we attempted to develop knock down protocols for both host and symbiont gene expression. We show that fluorescently-labelled antisense molecules reach the *B. saltans* cytoplasm as well as intracellularly localized bacteria. However, antisense inhibition was not achieved. Upon antisense compound treatment, *B. saltans* cells proliferate excessively, changing the protein composition of the cells. We hypothesize that *B. saltans* (or the bacteria it feeds upon) consume antisense molecules and experience growth boost by a mechanism unrelated to antisense inhibition.

## Results

### Identifying carriers able to deliver antisense molecules to *B. saltans* and its intracellular symbiont

Initially, we determined the best delivery protocol to introduce antisense molecules into *B. saltans* and Cbv cells. We treated *B. saltans* cells with fluorophore-conjugated synthetic peptide nucleic acids (PNAs), as these molecules have been most thoroughly tested in intracellular bacteria. For intracellular delivery, we used two different protocols: incubation of *B. saltans* cells for 24h and electroporation of *B. saltans* with 50 μM of peptide nucleic acid molecule PNA00218_TMR (Table 1).

**Table 1.**
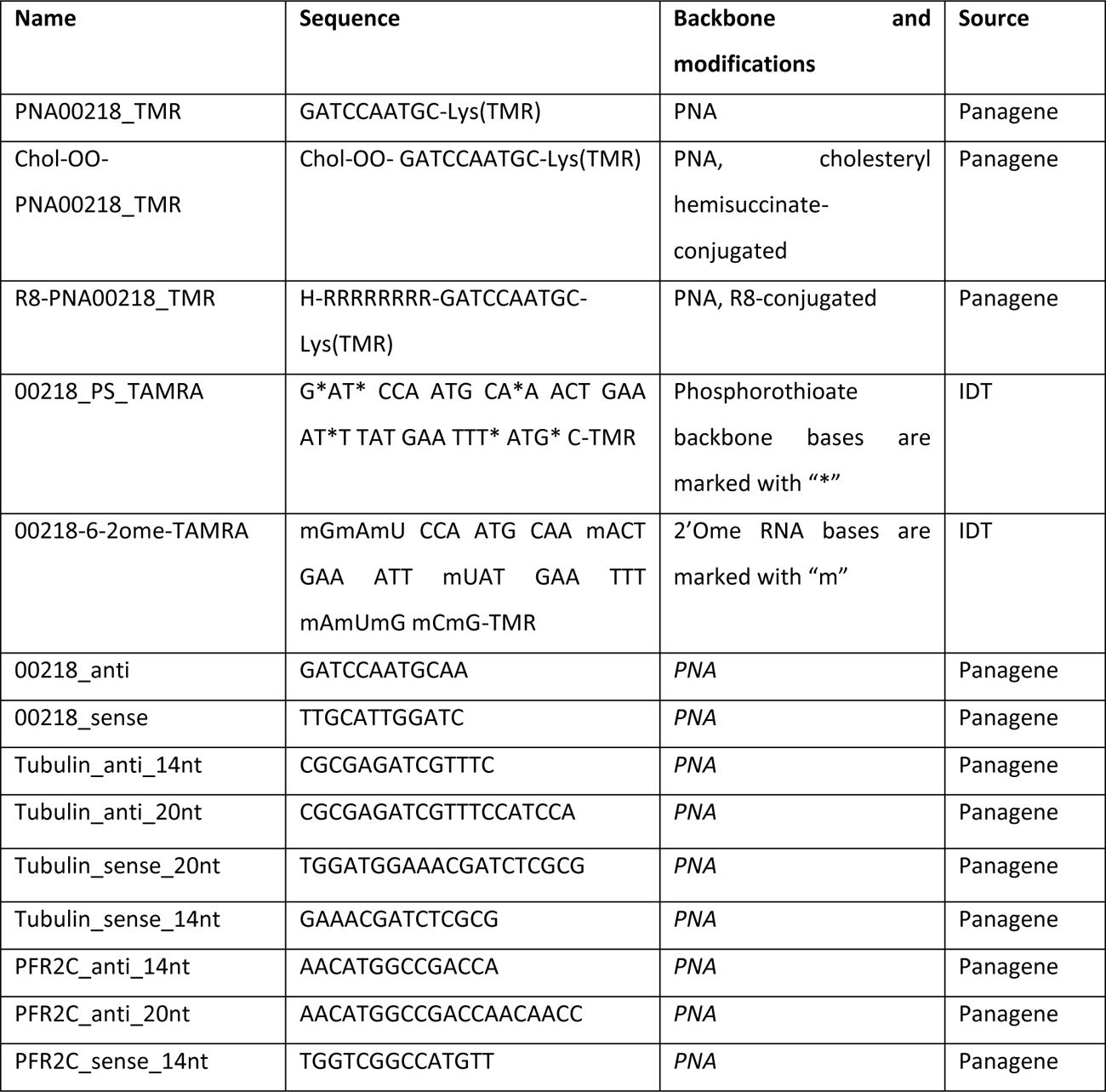
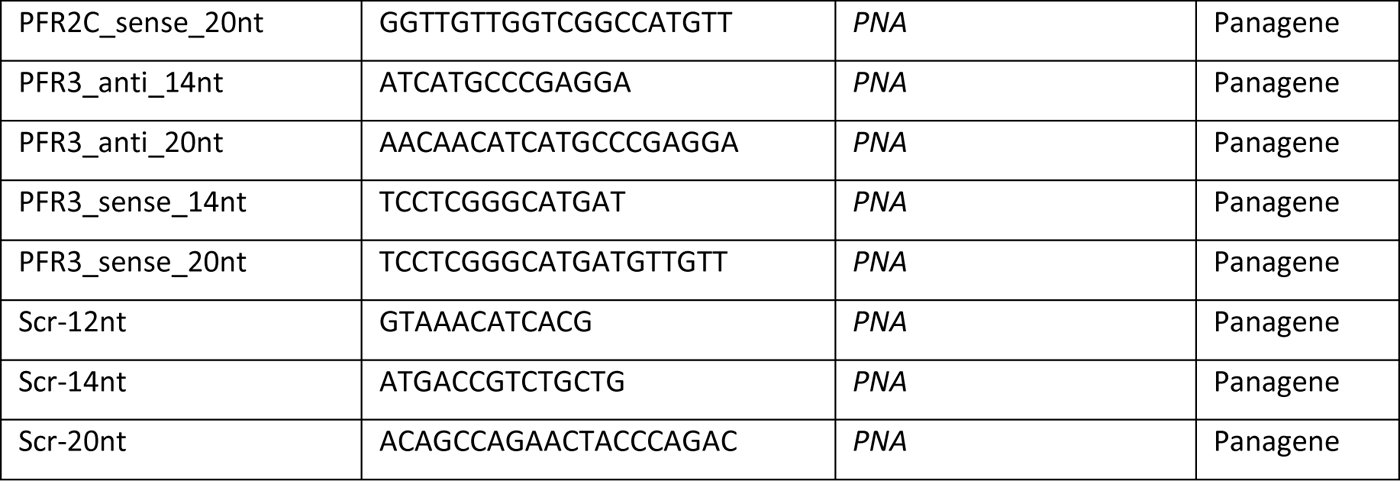
Oligonucleotide analogs designed and tested for B. saltans and its symbiont.

We imaged *B. saltans* cells after 24h to ascertain localization of the fluorescent antisense molecule within hosts and symbionts (Fig. 1A). In our experiment, incubation resulted in much higher fluorescence intensity within the *B. saltans* cells compared to electroporation (Fig. 1A, 1B, S1). This impact might be related to the fact that incubated cells are in contact with antisense molecules for 24hr while the electroporation protocol allows for just a very brief contact with antisense molecule (during electroporation).

**Figure 1.**
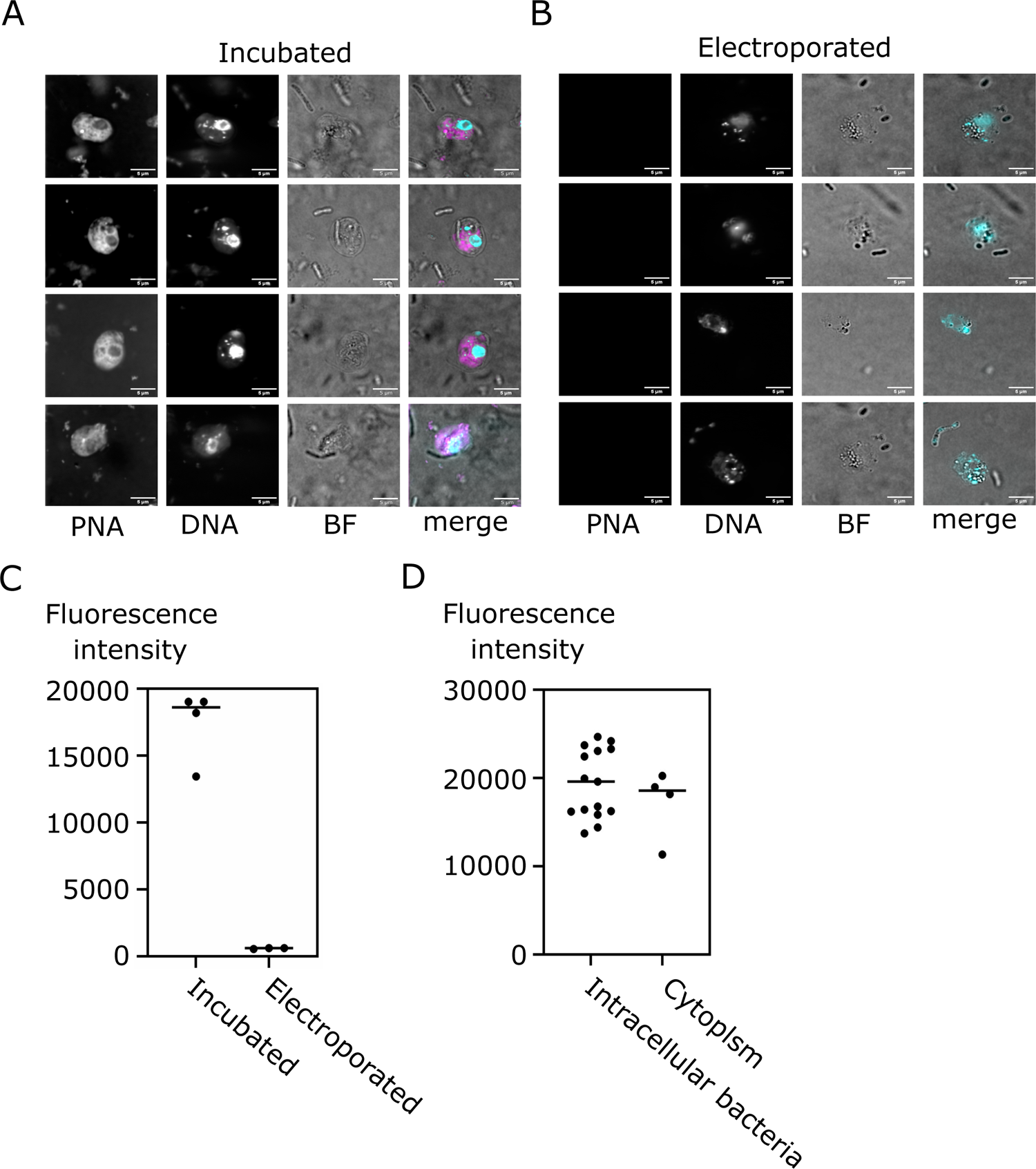
Incubation is more efficient than electroporation in delivering fluorescent PNAs to *B. saltans* and Cbv. Fluorescent images of *B. saltans* A) incubated or B) electroporated with fluorescently labelled antisense molecules. In merged images DNA is in cyan, TMR in magenta. C) Quantification of fluorescence intensity in the cytoplasm of *B. saltans* cells depicted in Fig. 1A and B. Only images acquired with the same settings were used for quantification. D) Quantification of fluorescence intensity in the intracellular bacteria and the cytoplasm (excluding the bacteria) in *B. saltans* incubated with antisense molecules depicted in Fig. 1A. Imaging parameters are in Table S1.

Next, we determined whether PNA00218_TMR can enter bacterial cells within *B. saltans* cytoplasm. To this end, we quantified the fluorescence intensity within the intracellular bacteria of *B. saltans* and compared it to the fluorescence intensity of the cell cytoplasm (excluding bacteria and nucleus). Intracellular bacteria had slightly higher average fluorescence intensity (19,383 AU vs 17,192 AU) than the surrounding cell cytoplasm, meaning that antisense molecules can enter Cbv cells but do not disproportionately accumulate within the symbiont.

In the process of optimizing the delivery we also assessed other antisense molecules chemistries, including PNAs attached to peptide and lipid carriers, 2’ome RNAs and phosphothiorate oligos (Table 1). However, they all consistently yielded weaker intracellular fluorescence by both incubation and electroporation (Fig. S2) in *B. saltans* cells. Incubation of *B. saltans* with PNAs became our protocol of choice for subsequent *B. saltans* and Cbv targeting experiments.

### Antisense inhibition of host and symbiont genes

To achieve antisense inhibition of *B. saltans* and intracellular symbiont, we designed antisense oligonucleotides spanning start codon and ribosome binding site of three host and two symbiont-encoded genes (18). The choice was dictated by the availability of specific antibodies for the host, while for the symbiont, we targeted proteins with readily predictable function. In *B. saltans*, our targets were PFR2C and PFR3 (paraflagellar rod protein isoforms, putatively involved in *B. saltans* feeding (24) and tubulin. For Cbv we used an antitoxin from one of the putative toxin-antitoxin operons (CPBP_00218 gene) and a putative cell division protein ftsZ. Bacterial antisense targeting is best achieved with 10 nucleotide-long antisense molecules (18). For *B. saltans*, genome complexity precludes the design of such short yet specific antisense molecules. Thus, we focused on 14 and 20 nucleotide antisense molecules. For all experiments, we have also used scrambled non-targeting oligos as a control.

First, we treated *B. saltans* cells with antisense PNAs for 24 and 48h designed to silence PFR2C and PFR3 (together for Western blot, as our antibody detects both protein isoforms, and separately for qPCR as we can detect each mRNA/cDNA separately) and tubulin (Fig. 2). We did not find evidence of antisense inhibition either by Western blot or by qPCR against targeted transcripts. Although protein samples were quantified and normalized for input, we observed stochasticity in PFR and Tubulin protein levels. Therefore, we have additionally confirmed that our Western blots were quantitative (Fig. S3), and that the differences in protein levels visible in Fig. 2A reflect *B. saltans* biology.

**Figure 2.**
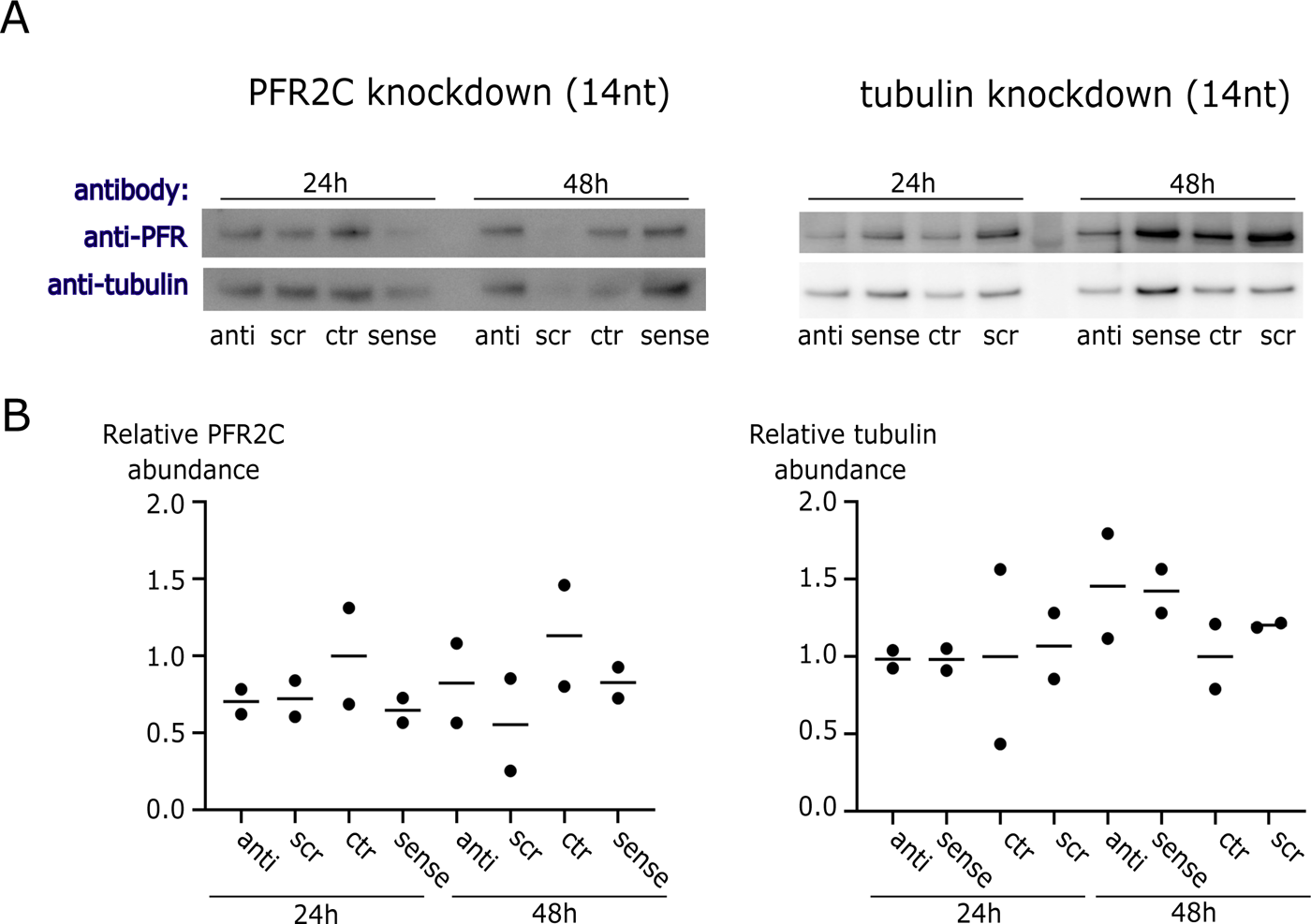
Antisense inhibition is not effective in *B. saltans*. A) Western blot with antibodies against PFR (L8C4) and tubulin (KMX-1) on whole *B. saltans* protein extract. *B. saltans* cells were treated with antisense molecules against PFR2C+PFR3 (the PFR isoforms recognized by the L8C4 antibody) and tubulin. Antisense, sense, scrambled and no antisense oligonucleotides were incubated with *B. saltans* cells for either 24 or 48h. B) Quantitative PCR for transcript abundance of genes targeted by antisense inhibition against PFR2C+PFR3 (left) or tubulin (right).

Due to the lack of specific antisense effect on *B. saltans* cells and the stochastic *B. saltans* protein abundance (especially in loading controls in the above experiments (Fig. 2)), we checked whether cell numbers of *B. saltans* were altered upon antisense PNA treatment (Fig. 3). To our surprise, *B. saltans* proliferated massively in culture upon antisense PNA addition. Although the strength of the effect depended on the oligonucleotide length and sequence, it was universal for the *B. saltans* and Cbv-targeting and non-targeting PNAs.

**Figure 3.**
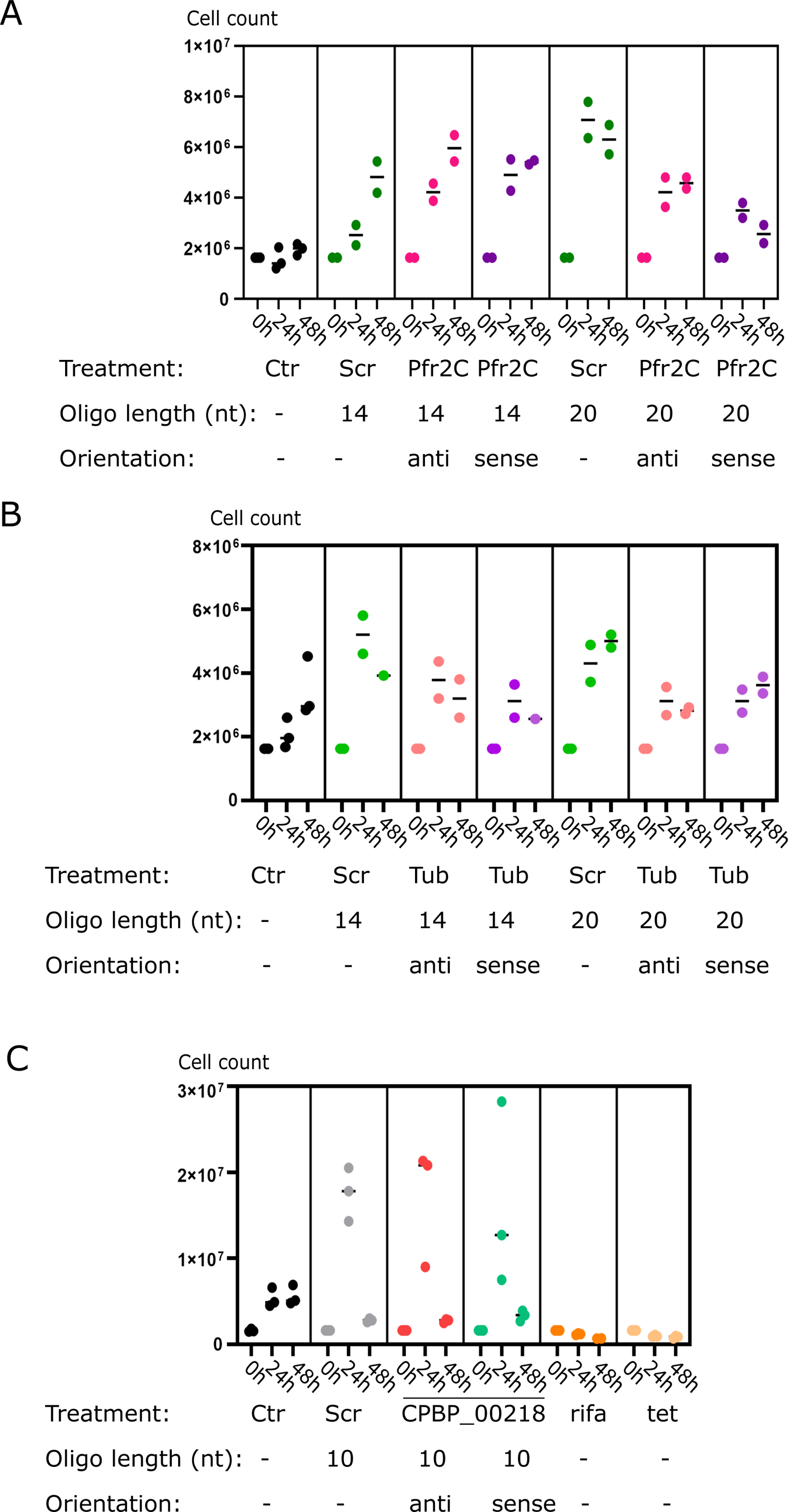
*B. saltans* proliferates massively when exposed to peptide nucleic acid molecules. Cell counts of *B. saltans* treated with antisense molecules against A) PFR2C+PFR3, B) Tubulin, C) Cbv gene CPBP_00218. In all cases *B. saltans* exposed to peptide nucleic acids reaches higher cell numbers. Rifa and tet are rifampicin and tetracycline, the controls used to see loss of *B. saltans* viability which we expected with CPBP_00218 gene knock down.

## Discussion

The study of obligate symbioses presents significant challenges for genetic analysis, particularly when dealing with non-model species and unculturable symbionts. We focused on the kinetoplastid host, *B. saltans*, and its bacterial symbiont Cbv, an association which exemplifies these challenges. Our aim was to develop tools for the analysis of gene function within this symbiosis using antisense inhibition. Our results demonstrated that antisense inhibition is not feasible in *B. saltans* and its symbiont under the conditions tested. Despite successful delivery of fluorescently-labelled antisense molecules into the *B. saltans* cells and into their intracellular bacteria, we did not achieve the anticipated knock-down of gene expression. Instead, we observed a surprising proliferation of *B. saltans* upon treatment with the antisense constructs. We have two potential explanations for this effect. First, that *B. saltans* itself may derive nutrition from the antisense molecules. Second, *Klebsiella* – the prey of *B. saltans* – may derive nutrition, indirectly fuelling *B. saltans* proliferation.

PNAs are synthetic mimics of DNA in which the deoxyribose phosphate backbone is replaced by a pseudo-peptide polymer to which the nucleobases (purines and pyrimidines) are linked (17, 25). Repetitive units of N-(2-aminoethyl) glycine constituting the backbone of PNAs have been shown to be from 30x to 1000x more resistant to different proteases than the peptides of the same length. However, they are not entirely resistant to proteolysis (26). *Bodo saltans* has a particularly high number of genes involved in the breakdown of peptides, and bacterial amino acids are the major source of energy, carbon, and nitrogen for this eukaryote (27). Additionally, *B. saltans* is a purine auxotroph (27). Therefore, PNAs in the medium might constitute an easy source of essential nutrients for *B. saltans*, promoting its growth.

Moreover, *B. saltans* feeds on bacteria – *K. pneumoniae* - in our laboratory cultures. These bacteria do not seem to quench the antisense molecules (Fig. 1A & B, note the extracellular bacteria in the bright field images). Nevertheless, they might be excreting proteases to the medium (28), impeding the delivery of full-length PNAs to the *B. saltans* and Cbv, even when the fluorophore is internalized. Additional nutrients produced in the process might feed *K. pneumoniae* or *B. saltans*, inducing proliferation of the latter.

The inability to achieve antisense inhibition in *B. saltans* and Cbv suggests that this method, while effective in other systems such as trypanosomes and Cbv-related bacteria (14–16, 19, 20, 29), may not be universally applicable across different species and symbiotic contexts. The unique metabolic and cellular processes of *B. saltans* or its microbiome might interfere with the antisense mechanism, necessitating alternative approaches for manipulation. Other gene targeting techniques such as homologous recombination and CRISPR/Cas9 are currently being established for *B. saltans* (23, 24), and further optimization might make them available for its intracellular symbionts as well. In the future, they will enable thorough investigation of the molecular basis of the symbiotic relationship between *B. saltans* and Cbv, including the role of the mysterious Bacterial Polymorphic Toxin Systems (PTS) in maintaining this obligate symbiosis.

Although our attempt to use antisense inhibition in *B. saltans* and its symbiont Cbv was unsuccessful, our study provides important insights and protocols that contribute to the broader effort of understanding and manipulating gene function in complex symbiotic systems. We developed electroporation, incubation, live imaging, image analysis, and Western blotting protocols, and identified a set of antibodies cross-reactive with *B. saltans*. These will strengthen our future efforts to identify genetic underpinnings of the obligate *B. saltans* – Cbv symbiosis.

## Materials and methods

### *B. saltans* culture

*B. saltans* was cultured in a cerophyl-based medium enriched with 3.5 mM sodium phosphate dibasic (Na_2_HPO_4_) (24). Cultures were incubated at 22°C in T25 tissue culture flasks containing 20 ml of media bacterized with *Klebsiella pneumoniae subsp. Pneumoniae* (ATCC® 700831). Three to 4 day old cultures were used for experiments.

### Antisense molecules

Antisense peptide nucleic acids were synthetised by Panagene, South Korea. Other oligonucleotides were synthesised by IDT. All antisense oligonucleotides were ordered with HPLC purification, >90% purity.

### Incubation of *B. saltans* cells with antisense molecules

*B. saltans* cultures were filtered through 100 and 8 μm filters. Cells were harvested by centrifugation at 1200 × g for 12 mins at 19°C, washed with 10-15 ml sterile filtered (SF) 1×PBS, and cells re-suspended in 5 ml SF 1×PBS. Single treatment cell aliquots containing 5×10^5^ cells were centrifuged again, and PBS replaced with SF cerophyll medium. Antisense molecules were added in a final concentration of 50 μM. After 24 h and 48 h *B. saltans* cells were harvested for Western blot and qPCR. The detailed protocol can be found here:dx.doi.org/10.17504/protocols.io.6qpvr8pqblmk/v1s.

### Electroporation of fluorescent antisense molecules into *B. saltans* and live imaging

We electroporated antisense molecules to *B. saltans* cells according to the published protocol (https://www.protocols.io/view/electroporation-of-fluorescent-antisense-molecules-e6nvwk8e2vmk/v1). To prepare *B. saltans* cells for electroporation, the culture was first filtered through 100 μm and 8 μm filters. The cells were then harvested by centrifugation at 1200 × g for 12 minutes at 19°C. Following this, the cells were washed with 10 ml PBS and centrifuged again under the same conditions. The cells were resuspended in 5 ml PBS, counted using a hemacytometer, and the volume containing 5×10^5^ cells was taken as recommended for the Neon transfection kit (Thermo Fisher Scientific) with a 10 μl tip. The cells were centrifuged once more at 1200 × g for 12 minutes at 19 °C. After removing the PBS, the cells were resuspended in electroporation buffer. An antisense molecule was then added to the *B. saltans* cells at a final concentration of 50 μM and mixed by pipetting. Finally, the mixture was aspirated into a Neon pipette and electroporated with a single pulse at 1800 V with a 10 ms pulse width.

### Imaging

In order to image *B. saltans* cells, we mixed incubated or electroporated cells with 1% low melting temperature agarose (Thermo Fisher Scientific) in 1:1 ratio and let it set for a few seconds in a well of a 96-well plate. We stained DNA with Hochest 33342 (Thermo Fisher Scientific) diluted 1:2000 in PBS by adding the solution to the agarose-embedded *B. saltans* and incubating it for 10 mins at room temperature (RT). We washed the agarose with PBS (2×5mins) at RT. We removed the agarose from the well with clean forceps, placed it on a microscope slide, added a drop of mounting medium (Vectashield, Vector Laboratories), and flattened the agarose using the coverslip. We proceeded with confocal imaging at the Centre for Cell Imaging using an LSM 880 Laser Scanning Confocal (Zeiss) equipped with a 63x (1.4 NA) objective, and parameters listed in Table S1. Because the bright field images were used to localize the *B*. saltans and for general reference only, the acquisition and display parameters were adjusted individually for each image. Images were analysed using custom Fiji (30) Script provided by Marie Held at the Centre for Cell Imaging, University of Liverpool. The protocol for analysis can be found here: dx.doi.org/10.17504/protocols.io.6qpvr8pqblmk/v1 and the script for automation of analysis can be found here: https://github.com/Marien-kaefer/General_Fiji_macros/tree/main/BodoSaltans

### Western blot

We performed Western blots according to the published protocol (https://www.protocols.io/view/protein-extraction-quantification-and-western-blot-5qpvobqodl4o/v2). Cells were harvested as above, and protein was extracted according to Newton *et al*. (31). Subsequently, protein was quantified with Pierce™ BCA Protein Assay Kit (Thermo Fisher Scientific) according to the manufacturer’s instructions. All concentrations were normalized to those of the most diluted samples. Samples were reduced, separated on polyacrylamide gels (Bolt Bis-Tris Plus, Thermo Fisher Scientific) and then transferred to PVDF membranes using a mini blot module (Thermo Fisher Scientific). The membranes were blocked for 30 minutes at room temperature in 5% skimmed milk diluted in wash buffer (PBS with 0.1% Tween 20). Subsequently, the membranes were incubated overnight at 4°C with primary antibodies (Thermo Fisher Scientific, diluted 1:100 in 5% milk in wash buffer, Table 3). After incubation, the membranes were washed three times for 10 minutes each with wash buffer. They were then incubated for 1 hour at room temperature with Horseradish Peroxidase (HRP)-labelled secondary antibodies (diluted 1:5000 in 5% milk in wash buffer). The membranes were washed again three times for 10 minutes each with wash buffer. Finally, the signal was developed using an HRP substrate (e.g., Immobilon western kit, Merck) and scanned for chemiluminescent signal using an imaging system (e.g., ImageQuant LAS4000 or Bio-Rad Chemidoc). The exposure time was chosen for each membrane separately to the highest value possible while avoiding saturated pixels, and results were saved as 16 bit images.

**Table 3.**
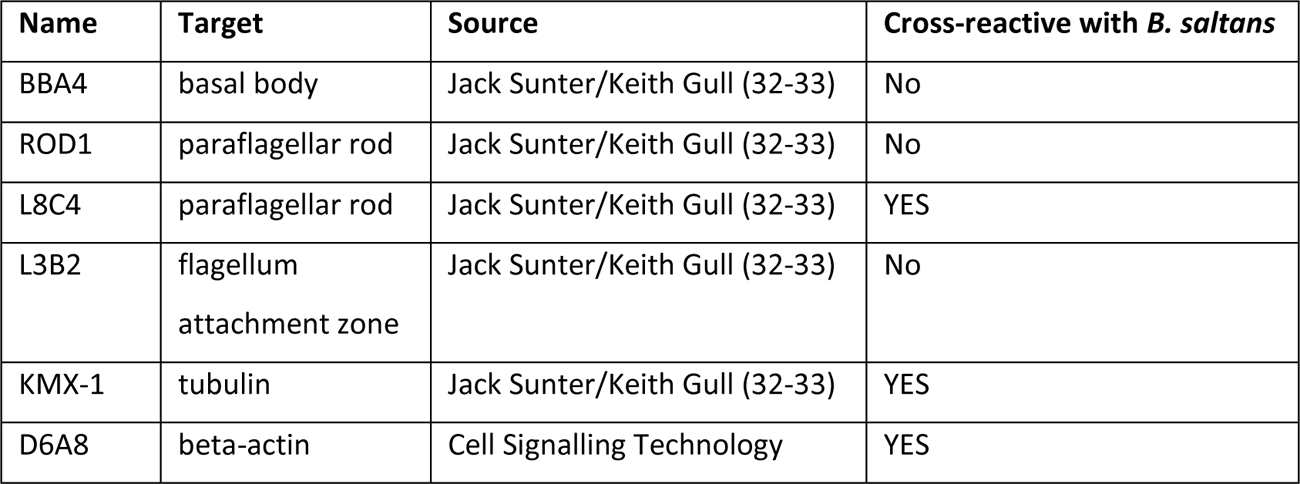
Antibodies tested for cross reactivity with B. saltans.

### RNA extraction, cDNA synthesis and qPCR

*B. saltans* cells were homogenized in Trizol Reagent (Invitrogen) and RNA extracted using a Direct-zol®RNA MiniPrep kit (Zymo Research) including a DNAse digestion step according to the manufacturer’s instructions. RNA was eluted in 25 μl DNase/RNase-Free Water (Zymo Research) and the concentration determined using a NanoDrop Spectrophotometer. cDNA was prepared from 1 μg of total RNA using Random Primers and M-MLV Reverse Transcriptase (both Promega). Primers were allowed to bind to template RNA for 5 minutes at 70°C, followed by 25°C for 10 minutes before M-MLV was added and incubated at 37°C for 60 minutes then 80°C for 10 minutes.

Real-time qPCRs were carried out in a LightCycler® 480 Instrument II (Roche) with the Power SYBR® Green PCR Master Mix (ThermoFisher Scientific). Each reaction contained 6 μl of SYBR® Green PCR Master Mix, 0.5 μl of each primer solution at 3.6 μM and 5 μl of diluted DNA. Each plate contained three technical replicates of every sample for each set of primers. Primers used are listed in Table 4. Relative expression levels were calculated by the Pfaffl method (34), with PFR2C relative to tubulin and tubulin relative to actin.

**Table 4.**
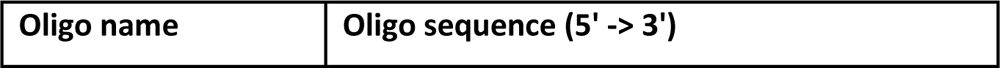
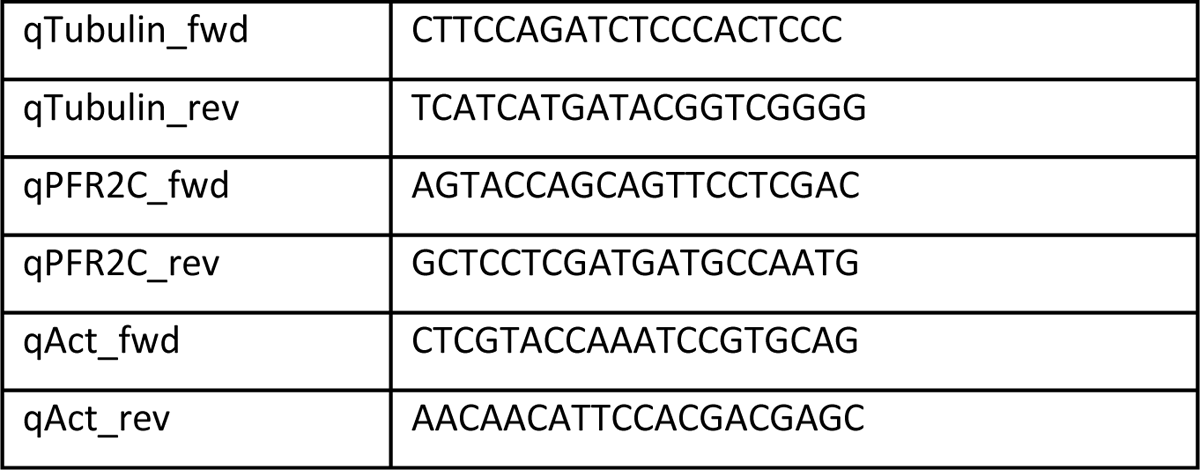
Oligonucleotide primers used for qPCR.

## Acknowledgements

We thank Dr. Samriti Middha for help with establishing *B. saltans* cultures. We are grateful to Dr. Jack Sunter (Oxford Brooks) and Prof. Keith Gull (Oxford University) for monoclonal antibodies against trypanosome proteins. We thank the Centre for Cell Imaging (CCI) at the University of Liverpool for the assistance with live *B. saltans* imaging. Zeiss 880 BioAFM at the CCI was funded by BBSRC grant number BB/M012441/1.

This work was funded by Gordon and Betty Moore Foundation’s Symbiosis in Aquatic Systems Initiative, Grant ID: #9357 (https://doi.org/10.37807/GBMF9357), awarded to GH, EC and AJ.

## Supplementary figure legends

**Figure S1.**
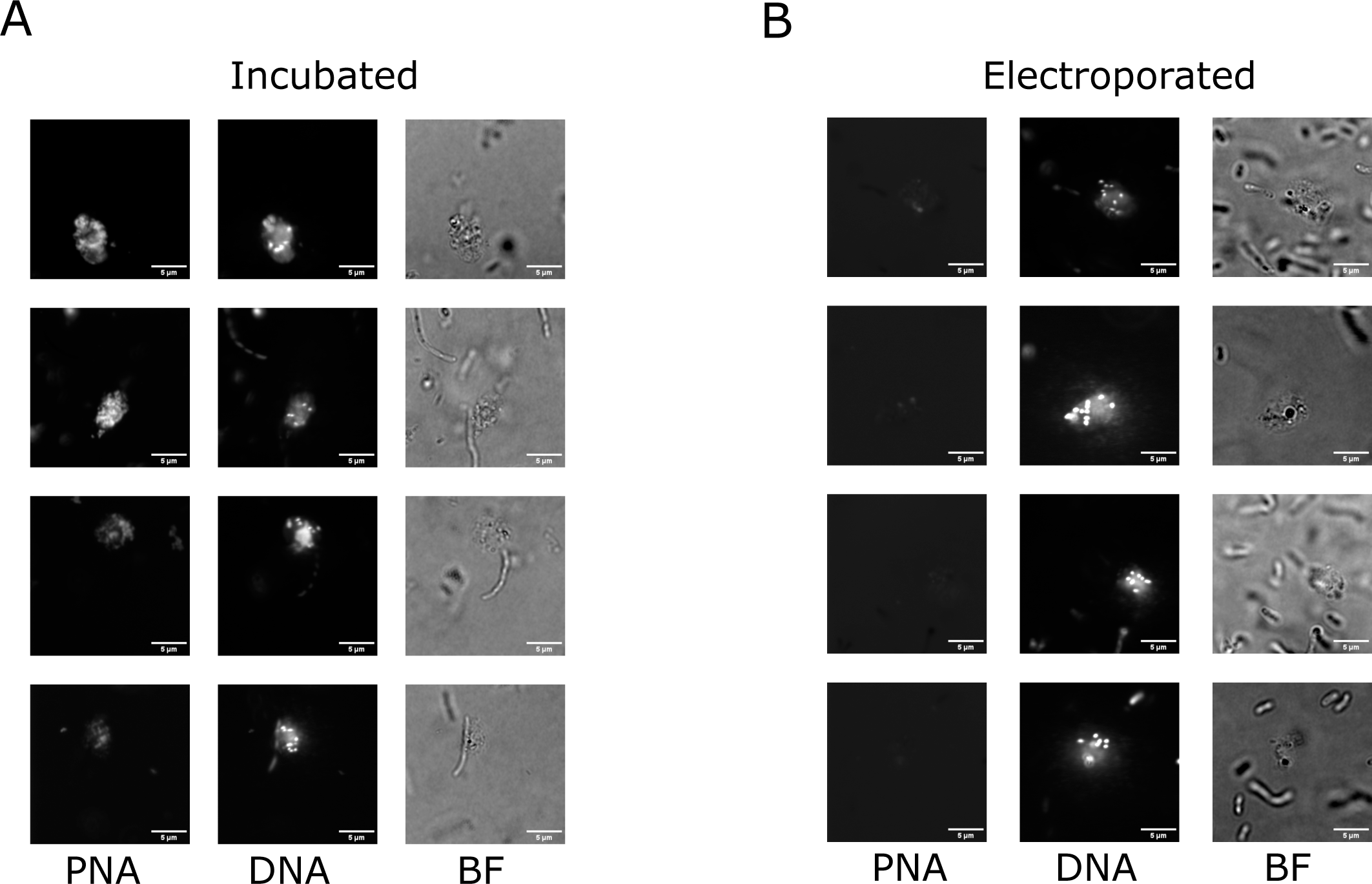
Incubation is more efficient than electroporation in delivering fluorescent PNAs *B. saltans* and Cbv. Fluorescent images of *B. saltans* A) incubated or B) electroporated with fluorescently labelled antisense molecules. This image was acquired with different TMR channel settings than the Fig. 1. Imaging parameters are in Table S1.

**Figure S2.**
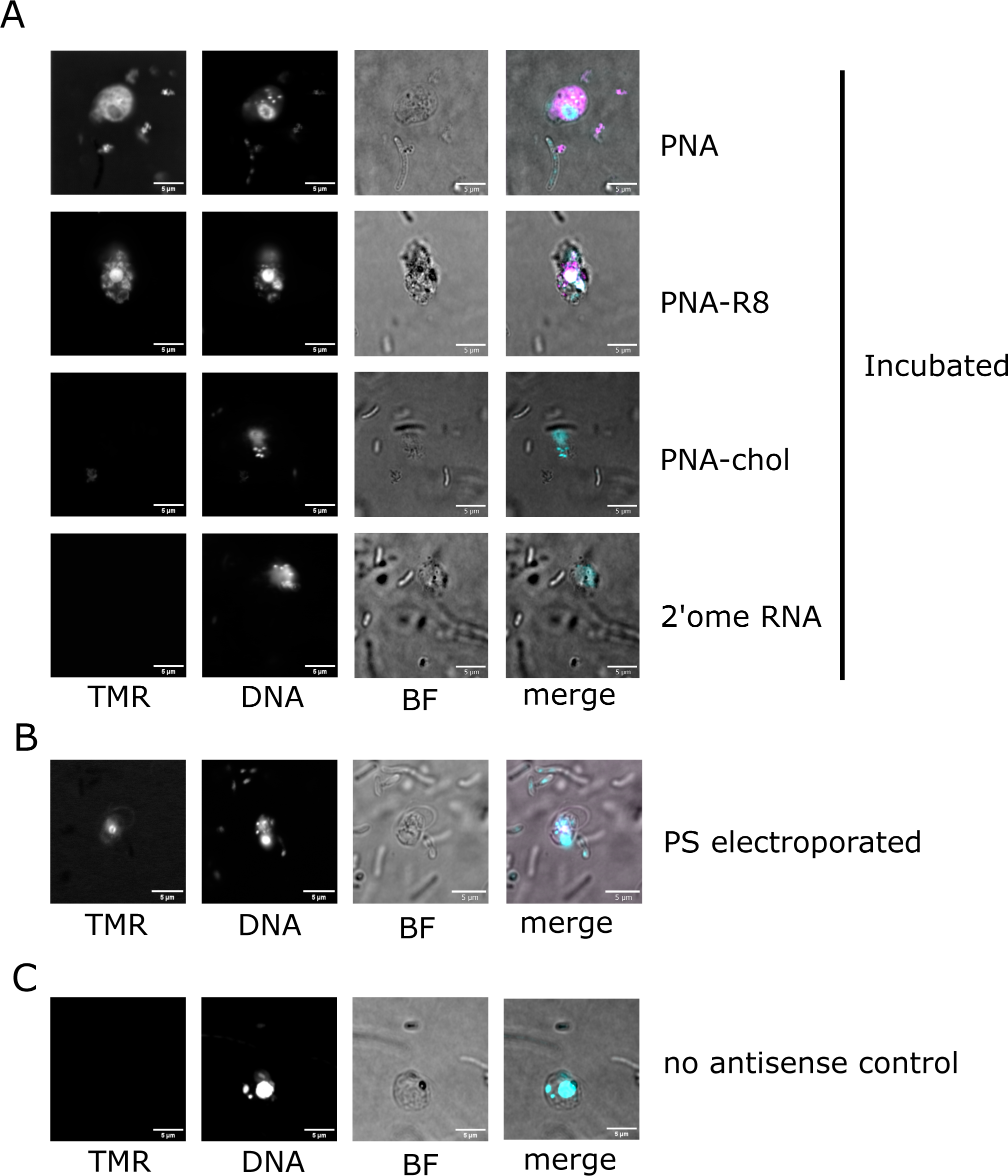
Untagged PNAs incubated with *B. saltans* are the most effective in entering microeukaryote cells. A) PNAs, PNAs tagged with a cell penetrating peptide R8 or cholesteryl chemisuccinate, and 2’ome RNA were incubated with the cells. B) Phosphothoreate oligo was electroporated into *B. saltans* and imaged using 3x zoom. This image was cropped for display but cannot be directly compared with the other images. C) No antisense molecule control to visualize potential *B. saltans* autofluorescence. In merged images DNA is in cyan, TMR in magenta. Sequences and modifications of each of the molecules are listed in Table 1. Imaging parameters are in Table S1.

**Figure S3.**
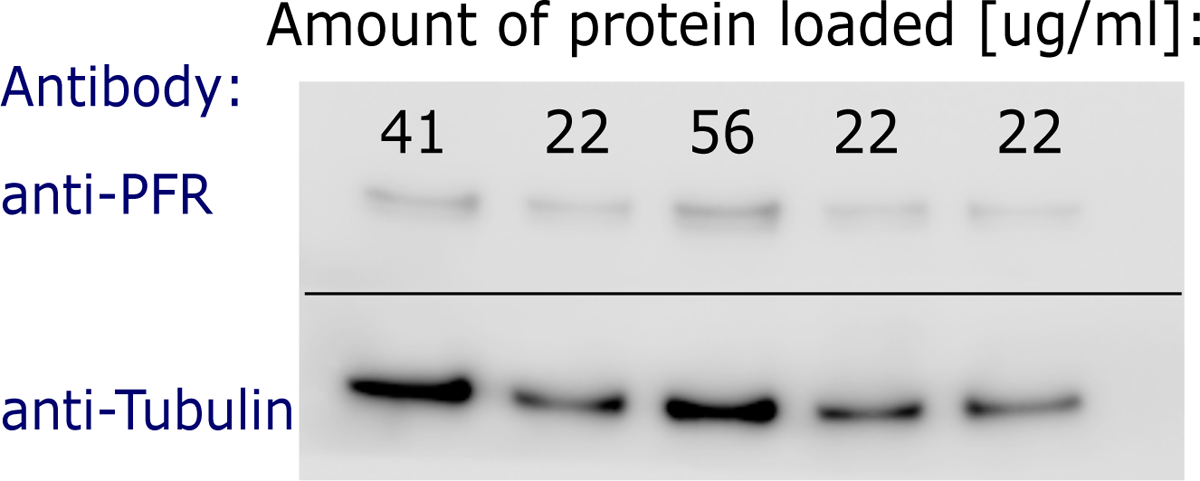
Western blot is a quantitative method for *B. saltans*. Western blot with antibodies against PFR (L8C4) and tubulin (KMX-1) were used to probe a membrane with different quantities of whole *B. saltans* protein extract.

