## Supplementary material for "Development of antisense tools to study *Bodo saltans* and its intracellular symbiont": Table S1

Fugure 1

| Treatment | FileName | Fluor | Ranges | PixelTime ( | Gain | Averaging | DigitalGain | Offset | Attenuatio | PinholeDiameter (m) | FileName | Fluor | Ranges | PixelTime ( | Gain | Averaging | DigitalGain | Offset | Attenuatio | PinholeDiameter (m) |
|---|---|---|---|---|---|---|---|---|---|---|---|---|---|---|---|---|---|---|---|---|
| Electroporated PNA | live dot electroporated 2.czi_metadata.xml | TMR | 564.25999999999999-665.23000000000025 | 2.67E-05 | 507.9828 | 16 | 1 | 0 | 0.114872 | 4.50E-05 | live dot electroporated 2.czi_metadata.xml | DAPI | 409.44999999999999-577.82000000000039 | 2.67E-05 | 700 | 16 | 1 | 0 | 0.642171 | 4.75E-05 |
| Electroporated PNA | live dot electroporated 3.czi_metadata.xml | TMR | 564.25999999999999-665.23000000000025 | 2.67E-05 | 507.9828 | 16 | 1 | 0 | 0.114872 | 4.50E-05 | live dot electroporated 3.czi_metadata.xml | DAPI | 409.44999999999999-577.82000000000039 | 2.67E-05 | 700 | 16 | 1 | 0 | 0.642171 | 4.75E-05 |
| Electroporated PNA | live dot electroporated 4.czi_metadata.xml | TMR | 564.25999999999999-665.23000000000025 | 2.67E-05 | 507.9828 | 16 | 1 | 0 | 0.114872 | 4.50E-05 | live dot electroporated 4.czi_metadata.xml | DAPI | 409.44999999999999-577.82000000000039 | 2.67E-05 | 700 | 16 | 1 | 0 | 0.642171 | 4.75E-05 |
| Electroporated PNA | live dot electroporated 6.czi_metadata.xml | TMR | 564.25999999999999-665.23000000000025 | 2.67E-05 | 508 | 16 | 1 | -1.14E-12 | 0.115 | 4.50E-05 | live dot electroporated 6.czi_metadata.xml | DAPI | 409.44999999999999-577.82000000000039 | 2.67E-05 | 535 | 16 | 1 | 0 | 0.642171 | 4.75E-05 |
| Incubated PNA | 13 dot incubated zoom6 speed4 obj63 .czi_metadata.xml | TMR | 564.25999999999999-665.23000000000025 | 2.67E-05 | 507.9828 | 16 | 1 | 0 | 0.114872 | 4.50E-05 | 13 dot incubated zoom6 speed4 obj63 .czi_metadata.xml | DAPI | 409.44999999999999-577.82000000000039 | 2.67E-05 | 700 | 16 | 1 | 0 | 0.642171 | 4.75E-05 |
| Incubated PNA | 18 dot incubated zoom6 speed4 obj63 .czi_metadata.xml | TMR | 564.25999999999999-665.23000000000025 | 2.67E-05 | 508 | 16 | 1 | 0 | 0.114872 | 4.50E-05 | 18 dot incubated zoom6 speed4 obj63 .czi_metadata.xml | DAPI | 409.44999999999999-577.82000000000039 | 2.67E-05 | 700 | 16 | 1 | 0 | 0.642171 | 4.75E-05 |
| Incubated PNA | 16 dot incubated zoom6 speed4 obj63 .czi_metadata.xml | TMR | 564.25999999999999-665.23000000000025 | 2.67E-05 | 508 | 16 | 1 | 0 | 0.114872 | 4.50E-05 | 16 dot incubated zoom6 speed4 obj63 .czi_metadata.xml | DAPI | 409.44999999999999-577.82000000000039 | 2.67E-05 | 700 | 16 | 1 | 0 | 0.642171 | 4.75E-05 |
| Incubated PNA | 17 dot incubated zoom6 speed4 obj63 .czi_metadata.xml | TMR | 564.25999999999999-665.23000000000025 | 2.67E-05 | 508 | 16 | 1 | 0 | 0.114872 | 4.50E-05 | 17 dot incubated zoom6 speed4 obj63 .czi_metadata.xml | DAPI | 409.44999999999999-577.82000000000039 | 2.67E-05 | 700 | 16 | 1 | 0 | 0.642171 | 4.75E-05 |

Figure S1

| Treatment | FileName | Fluor | Ranges | PixelTime ( | Gain | Averaging | DigitalGain | Offset | Attenuatio | PinholeDiameter (m) | FileName | Fluor | Ranges | PixelTime ( | Gain | Averaging | DigitalGain | Offset | Attenuatio | PinholeDiameter (m) |
|---|---|---|---|---|---|---|---|---|---|---|---|---|---|---|---|---|---|---|---|---|
| Electroporated PNA | 1st slide dot electroporated tamra 508 gain 3.czi_metadata | TMR | 564.25999999999999-665.23000000000025 | 2.67E-05 | 508 | 16 | 1 | -1.14E-12 | 0.115 | 4.50E-05 | 1st slide dot electroporated tamra 508 gain 3.czi_metadata.xm | DAPI | 409.44999999999999-577.82000000000039 | 2.67E-05 | 672.2275 | 16 | 1 | 0 | 0.642171 | 4.75E-05 |
| Electroporated PNA | 1st slide dot electroporated tamra 508 gain 5.czi_metadata | TMR | 564.25999999999999-665.23000000000025 | 2.67E-05 | 508 | 16 | 1 | -1.14E-12 | 0.115 | 4.50E-05 | 1st slide dot electroporated tamra 508 gain 5.czi_metadata.xm | DAPI | 409.44999999999999-577.82000000000039 | 2.67E-05 | 626.2189 | 16 | 1 | 0 | 0.642171 | 4.75E-05 |
| Electroporated PNA | 1st slide dot electroporated tamra 508 gain 7.czi_metadata | TMR | 564.25999999999999-665.23000000000025 | 2.67E-05 | 508 | 16 | 1 | -1.14E-12 | 0.115 | 4.50E-05 | 1st slide dot electroporated tamra 508 gain 7.czi_metadata.xm | DAPI | 409.44999999999999-577.82000000000039 | 2.67E-05 | 580.2103 | 16 | 1 | 0 | 0.642171 | 4.75E-05 |
| Electroporated PNA | 1st slide dot electroporated tamra 508 gain 8.czi_metadata | TMR | 564.25999999999999-665.23000000000025 | 2.67E-05 | 508 | 16 | 1 | -1.14E-12 | 0.115 | 4.50E-05 | 1st slide dot electroporated tamra 508 gain 8.czi_metadata.xm | DAPI | 409.44999999999999-577.82000000000039 | 2.67E-05 | 580.2103 | 16 | 1 | 0 | 0.642171 | 4.75E-05 |
| Incubated PNA | 2 incubated dot tamra 508.czi_metadata.xml | TMR | 564.25999999999999-665.23000000000025 | 2.67E-05 | 508 | 16 | 1 | -1.14E-12 | 0.115 | 4.50E-05 | 2 incubated dot tamra 508.czi_metadata.xml | DAPI | 409.44999999999999-577.82000000000039 | 2.67E-05 | 635.4206 | 16 | 1 | 0 | 0.642171 | 4.75E-05 |
| Incubated PNA | 4 incubated dot tamra 508.czi_metadata.xml | TMR | 564.25999999999999-665.23000000000025 | 2.67E-05 | 508 | 16 | 1 | -1.14E-12 | 0.115 | 4.50E-05 | 4 incubated dot tamra 508.czi_metadata.xml | DAPI | 409.44999999999999-577.82000000000039 | 2.67E-05 | 589.412 | 16 | 1 | 0 | 0.642171 | 4.75E-05 |
| Incubated PNA | 11 incubated dot tamra 508.czi_metadata.xml | TMR | 564.25999999999999-665.23000000000025 | 2.67E-05 | 508 | 16 | 1 | -1.14E-12 | 0.115 | 4.50E-05 | 11 incubated dot tamra 508.czi_metadata.xml | DAPI | 409.44999999999999-577.82000000000039 | 2.67E-05 | 736.6395 | 16 | 1 | 0 | 0.642171 | 4.75E-05 |
| Incubated PNA | 14 incubated dot tamra 508.czi_metadata.xml | TMR | 564.25999999999999-665.23000000000025 | 2.67E-05 | 508 | 16 | 1 | -1.14E-12 | 0.115 | 4.50E-05 | 14 incubated dot tamra 508.czi_metadata.xml | DAPI | 409.44999999999999-577.82000000000039 | 2.67E-05 | 653.824 | 16 | 1 | 0 | 0.642171 | 4.75E-05 |

| Fugure S2 | FileName | Fluor | Ranges | PixelTime ( | Gain | Averaging | DigitalGain | Offset | Attenuatio | PinholeDiameter (m) | FileName | Fluor | Ranges | PixelTime ( | Gain | Averaging | DigitalGain | Offset | Attenuatio | PinholeDiameter (m) |
|---|---|---|---|---|---|---|---|---|---|---|---|---|---|---|---|---|---|---|---|---|
| No antisense | 1 control electroporated without antisenseTamara 508.czi | TMR | 564.25999999999999-665.23000000000025 | 2.67E-05 | 508 | 16 | 1 | -1.14E-12 | 0.115 | 4.50E-05 | 1 control electroporated without antisenseTamara 508.czi_me | DAPI | 409.44999999999999-577.82000000000039 | 2.67E-05 | 690.6309 | 16 | 1 | 0 | 0.642171 | 4.75E-05 |
| No antisense | 2 control electroporated without antisenseTamara 508.czi | TMR | 564.25999999999999-665.23000000000025 | 2.67E-05 | 508 | 16 | 1 | -1.14E-12 | 0.115 | 4.50E-05 | 2 control electroporated without antisenseTamara 508.czi_me | DAPI | 409.44999999999999-577.82000000000039 | 2.67E-05 | 920.6738 | 16 | 1 | 0 | 0.642171 | 4.75E-05 |
| Incubated 2'Ome RNA | 3heart incubated 508 tamra.czi_metadata.xml | TMR | 564.25999999999999-665.23000000000025 | 2.67E-05 | 508 | 16 | 1 | -1.14E-12 | 0.115 | 4.50E-05 | 3heart incubated 508 tamra.czi_metadata.xml | DAPI | 409.44999999999999-577.82000000000039 | 2.67E-05 | 635.4206 | 16 | 1 | 0 | 0.642171 | 4.75E-05 |
| Incubated Chol-PNA | 6 triangle incubated tamra 508.czi_metadata.xml | TMR | 564.25999999999999-665.23000000000025 | 2.67E-05 | 508 | 16 | 1 | -1.14E-12 | 0.115 | 4.50E-05 | 6 triangle incubated tamra 508.czi_metadata.xml | DAPI | 409.44999999999999-577.82000000000039 | 2.67E-05 | 653.824 | 16 | 1 | 0 | 0.642171 | 4.75E-05 |
| Incubated R8-PNA | 7 tick incubated tamra 508.czi_metadata.xml | TMR | 564.25999999999999-665.23000000000025 | 2.67E-05 | 508 | 16 | 1 | -1.14E-12 | 0.115 | 4.50E-05 | 7 tick incubated tamra 508.czi_metadata.xml | DAPI | 409.44999999999999-577.82000000000039 | 2.67E-05 | 617.0172 | 16 | 1 | 0 | 0.642171 | 4.75E-05 |
| Incubated PNA | 7dot incubated zoom6 speed4 obj63.czi_metadata.xml | TMR | 564.25999999999999-665.23000000000025 | 2.67E-05 | 507.9828 | 16 | 1 | 0 | 0.114872 | 4.50E-05 | 7dot incubated zoom6 speed4 obj63.czi_metadata.xml | DAPI | 409.44999999999999-577.82000000000039 | 2.67E-05 | 700 | 16 | 1 | 0 | 0.642171 | 4.75E-05 |
| Incubated Phosphorothioa | 63x obj 3x zoom electroporated cross a.czi_metadata.xml | TMR | 564.25999999999999-665.23000000000025 | 3.27E-06 | 747.4292 | 16 | 1 | 0 | 0.065 | 6.57E-05 | 63x obj 3x zoom electroporated cross a.czi_metadata.xml | DAPI | 409.44999999999999-577.82000000000039 | 3.27E-06 | 778 | 16 | 1 | 0 | 0.033261 | 4.75E-05 |
